# Probing universal protein dynamics using residue-level Gibbs free energy

**DOI:** 10.1101/2020.09.30.320887

**Authors:** Jochem H. Smit, Srinath Krishnamurthy, Bindu Y. Srinivasu, Rinky Parakra, Spyridoula Karamanou, Anastassios Economou

## Abstract

Hydrogen Deuterium Exchange Mass Spectrometry is a powerful monitor of protein intrinsic dynamics, yet the interpretation, visualization and cross-comparison of HDX-MS datasets is challenging. Here we present PyHDX, an open-source python package and web server, that batch-extracts the universal quantity Gibbs free energy at residue level over multiple protein conditions and homologues. Δ*G* values relate to protein normal modes and together provide a universal measure of protein flexibility.

**Availability:** PyHDX source code is released under the MIT license and can be accessed on [GitHub](https://github.com/Jhsmit/PyHDX).

---

Intrinsic dynamics underlie protein function[1]. Hydrogen/deuterium exchange mass spectrometry (HDX-MS) is a powerful monitor of intrinsic dynamics at near residue resolution[2]. It identifies differentially dynamic regions because these undergo H/D-exchange at different rates. In typical ‘bottom-up’ HDX-MS (Fig. 1a)[3], proteins are D-labelled in deuterated buffers over timescales spanning several orders of magnitude[4]. Exchange is quenched to limit back-exchange to 30% (by reducing the intrinsic rate of D-exchange through lowered pH/temperature[5]), proteins are proteolyzed into multiple overlapping peptides that are analysed by liquid chromatography-MS. D-uptake on each peptide is calculated from mass changes between the deuterated (+1 mass unit) and undeuterated form[6].

**Figure 1:**
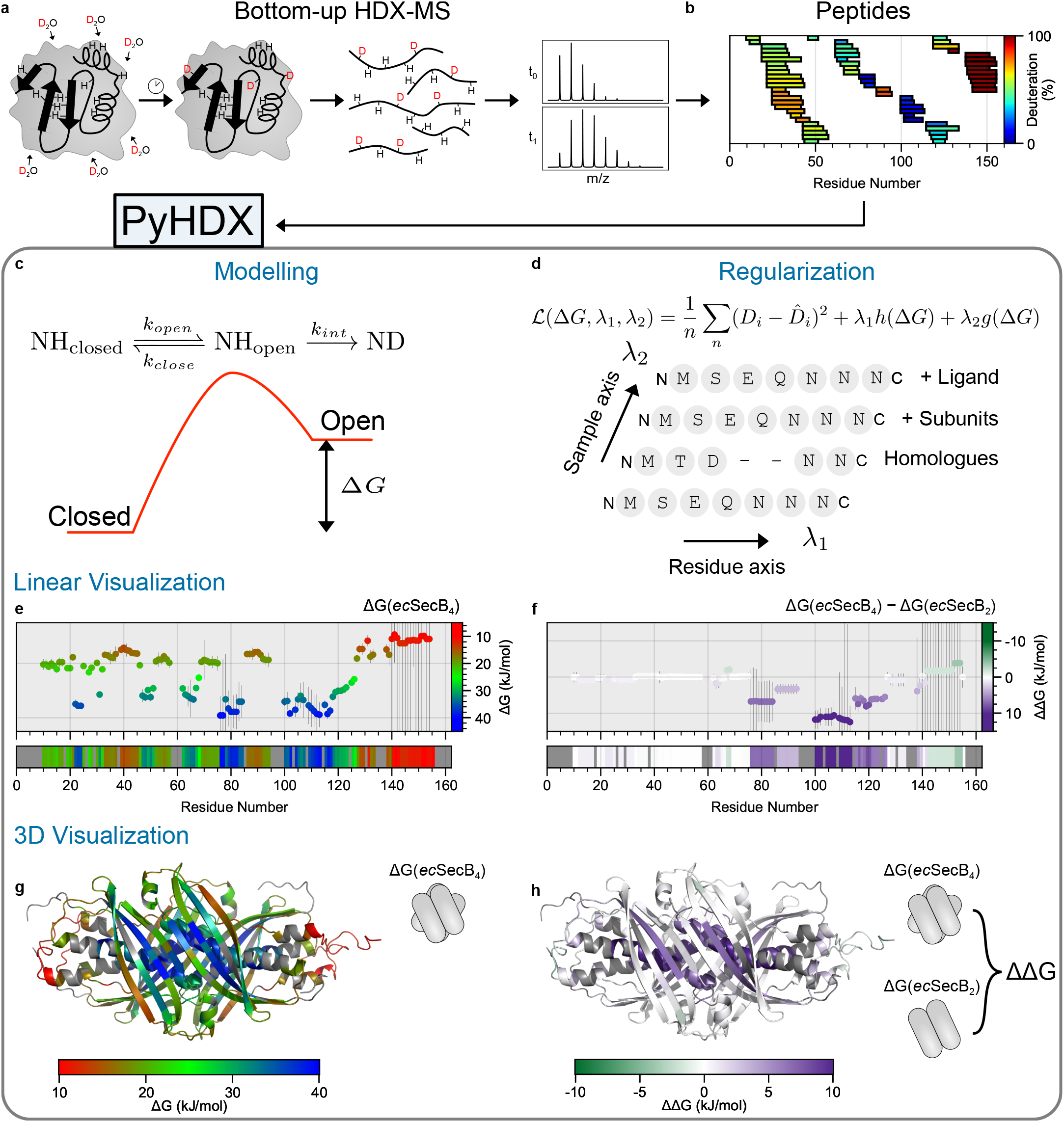
Pipeline of a bottom up HDX-MS experiment (top) analysed with PyHDX (bottom). **a**, HDX-MS experimental workflow consisting of deuterium exposure, quenching and pepsin digestion, and identification of peptides by LC/MS[7]. **b**, Overlapping peptides obtained as in a for *ec*SecB (UniProt P0AG86) showing a D-uptake heatmap relative to a fully deuterated control sample for all derivative peptides at t=30 s. **c**, Linderstrøm-Lang model[8, 9] of H/D exchange used in PyHDX to describe exchange in terms of Gibbs free energy (Δ*G*) between the closed non-exchanging and the open exchange-competent states. **d**, Two regularizers (*λ*_1_, *λ*_2_) are applied across two axes: the residue axis (*λ*_1_), minimizing variation in Δ*G* between consecutive residues, and the sample axis (*λ*_2_), minimizing variation in Δ*G* between residues along a set of HDX conditions (e.g. ligands, oligomeric state) or homologues. **e**, Output Δ*G* values per residue plotted against the linear sequence of *ec*SecB (top) coloured according to a gradient colour map (right). Residue colours are additionally shown as a linear bars (bottom), regions without peptide coverage are coloured grey. Error bars are covariances (see Methods). **f**, Differential dynamics between *ec*SecB_4_ and *ec*SecB_2_ shown as differences in Δ*G* of both states (ΔΔ*G*). Regions in purple are rigidifying in *ec*SecB_4_ compared to the dimeric state **g**, 3D structure of *ec*SecB_4_ coloured according to Δ*G*/residue from e. (PDB 5JTR[10], ligand removed) **h**, 3D structure of *ec*SecB_4_ coloured according to the per residue ΔΔ*G* from **f.**

D-uptake values are generally presented as heatmaps (Fig. 1b) or curves (Supplementary Fig. 1). These 2D slices of the full 3D dataset (time, deuteration, peptide; Supplementary Fig. 1), fail to capture the full breadth of the experimental information. It is desirable to display HDX-MS datasets as a single per residue value. For this, the overlap between peptides is exploited[2, 11–19] since HDX-MS yields deuteration values of each peptide *in toto* and not those of its individual amino acids. HDX-MS faces four remaining hurdles. Specifically, a) the large parameter space can render data analysis several hours-long b) experimental variations hamper differential dynamics between multiple datasets. c) Analysis tools often have complicated installation or commercial licensing. d) for HDX-MS-derived protein dynamics to have a wider impact and interface with orthogonal methods, output should yield universal quantities (e.g. Gibbs free energy). PyHDX addresses these issues and derives Gibbs free energy (Δ*G*), at residue level. Data is input as ‘HDX data’ tables in CSV format (peptide list, D-exposure time and D-uptake), submitted either as a single or multiple experiments. The full analysis and Gibbs free energy level classification of residues and visualization is accomplished in a web interface (Supplementary Fig. 2, Supplementary Video 1), within minutes and exported as text or a script to colour 3D structures in PyMOL (Schródinger, LLC). We used the commonly employed Linderstrøm-Lang model (Fig. 1c) for H/D exchange[8, 9]. In this model, backbone amides can be either in D-exchange-incompetent ‘closed state’, with amide hydrogens hydrogen-bonded in secondary structures, or in a D-exchange-competent ‘open state’, with compromised hydrogen bonds. The rate of D-exchange measured by HDX-MS allows calculation of Gibbs free energy differences (Δ*G*) between ‘open’/’closed’ states and the position of the equilibrium (i.e. protection factor; PF; popular in NMR studies[20]) through the expression Δ*G* = *RT* ln(*PF*). By assuming that the fractional population of NH_*open*_ is small and the rates of protein dynamics are faster than the intrinsic rate of H/D exchange (*k_int_*), an expression for D-uptake over time is obtained depending only on a single parameter per residue, thereby reducing complexity/degrees of freedom during fitting (see Methods). These approximations are generally true in the so called EX2 regime of H/D-exchange (i.e. *k_close_* ≫ *k_int_*, Fig. 1c)[21]. In EX1 (*k_close_* ≪ *k_int_*), PyHDX informs on differential dynamics qualitatively, as the approximations made break down introducing errors. To find Δ*G* values which best describe the data, we formulate a Lagrangian (cost function), composed of the Mean Square Error (MSE) and two regularizers (Lagrange Multipliers)[16, 17] (Fig. 1d), that constrain the possible solutions of Δ*G*. As amino acids of a peptide each have varying intrinsic H/D exchange rates that HDX-MS cannot determine directly, similar values of the Lagrangian can be satisfied with multiple combinations of Δ*G* assignment per amino acid, leading to non-identifiability[6]. To alleviate this, a regularizer *λ*_1_ acts along the primary sequence minimizing differences in Δ*G* between consecutive residues, unless experimental D-uptake values support such differences. HDX-MS informs on allostery by probing a protein’s dynamic response to external triggers, e.g. changes in oligomerization, mutations or ligands[22]. Such differential HDX-MS experiments[3, 23], compare D-uptake of reference and test states. Differential dynamics can be obtained by simply subtracting two datasets of PyHDX-derived single Δ*G*/residue. However, experimental variables (e.g. proteolysis, exchange timepoints and/or *k_int_*) can lead to artefactual “differences” between datasets. To alleviate this, a second regularizer (*λ*_2_) operates along the sample axis, minimizing differences between identical residues across datasets, unless experimental data support such a difference (Fig. 1d). We assessed PyHDX on the tetrameric *E. coli* chaperone SecB (*ec*SecB_4_)[24]. D-uptake was measured across six timepoints (10 sec-100 min; 30°C, pH_read_ 8; peptide heatmap (t=30 s; Fig. 1b); D-uptake time-kinetics curves for selected peptides; Supplementary Fig. 1). PyHDX-calculated Gibbs free energies for all residues (Fig. 1e; regularizer *λ*_1_ = 0.05). Setting *λ*_1_ sufficiently low allows the algorithm to extract features at high-resolution (Supplementary Fig. 3).). Covariances (see Methods) are shown as error bars, where a high covariance indicates that the protein’s flexibility in this regions lies outside of the range of Δ*G* values resolved by the experiment (as determined by temperature, pH and particularly timepoints). The obtained energies were classified into three regimes of relative flexibility: ‘rigid’ (40kJ mol^-1^, blue), ‘flexible’, (25 kJ mol^-1^, green) and ‘hyper-flexible’ (10 kJ mol^-1^, red), and assigned colours by linear interpolation between them. The resulting energy landscape was visualized onto the structure of *ec*SecB_4_ (Fig. 1g). Δ*G* values were also obtained for the dimeric mutant *ec*SecB_2_ (*λ*_1_ = 0.05, *λ*_2_ = 0.1) and values from the two states (ΔΔ*G*) were subtracted and coloured differently (10kJ mol^-1^, dark purple-increased rigidity; 0 kJ mol^-1^, white-no change; 10 kJ mol^-1^; dark green-increased dynamics). In *ec*SecB_4_, the internalized multimerization helix and the preceding *β*4 strand are most rigidified. ΔΔ*G* values of near zero indicate that data is insufficient to substantiate dynamics differences between two states. Specifically, in regions of extreme dynamics (e.g. highly disordered/rigid), experimental D-exchange timepoints must be chosen to adequately resolve differences if present. In the absence of such measurements, the regularizer *λ*_2_ ensures that ΔΔ*G* values in these regions show no change (e.g. *ec*SecB_4_ c-tail, residues 140-160, Supplementary Fig. 4) Next, we tested the dynamics of the 901-residue SecA (Fig. 2a), in 6 different biochemical contexts (monomer, dimer, ΔC-tail; +/-ADP)[25]. Despite the larger computational challenge (8883 peptide-timepoints; 5364 fit parameters), PyHDX converged to a solution within 5 minutes. PyHDX fits several states in parallel, at manageable computation times (e.g. 25 minutes to test 26 conditions, 23244 fit parameters; not shown). Δ*G* and ΔΔ*G* values were mapped onto linear maps of SecA_1_ or SecA_2_ or SecA-ΔC_2_ (Fig. 2a) and on the SecA_1_ structure (Figs. 2a and 2b). Nucleotide binding decreased dynamics/increased ΔΔ*G* mainly in helicase motifs[25, 26] (Fig 2c).

**Figure 2:**
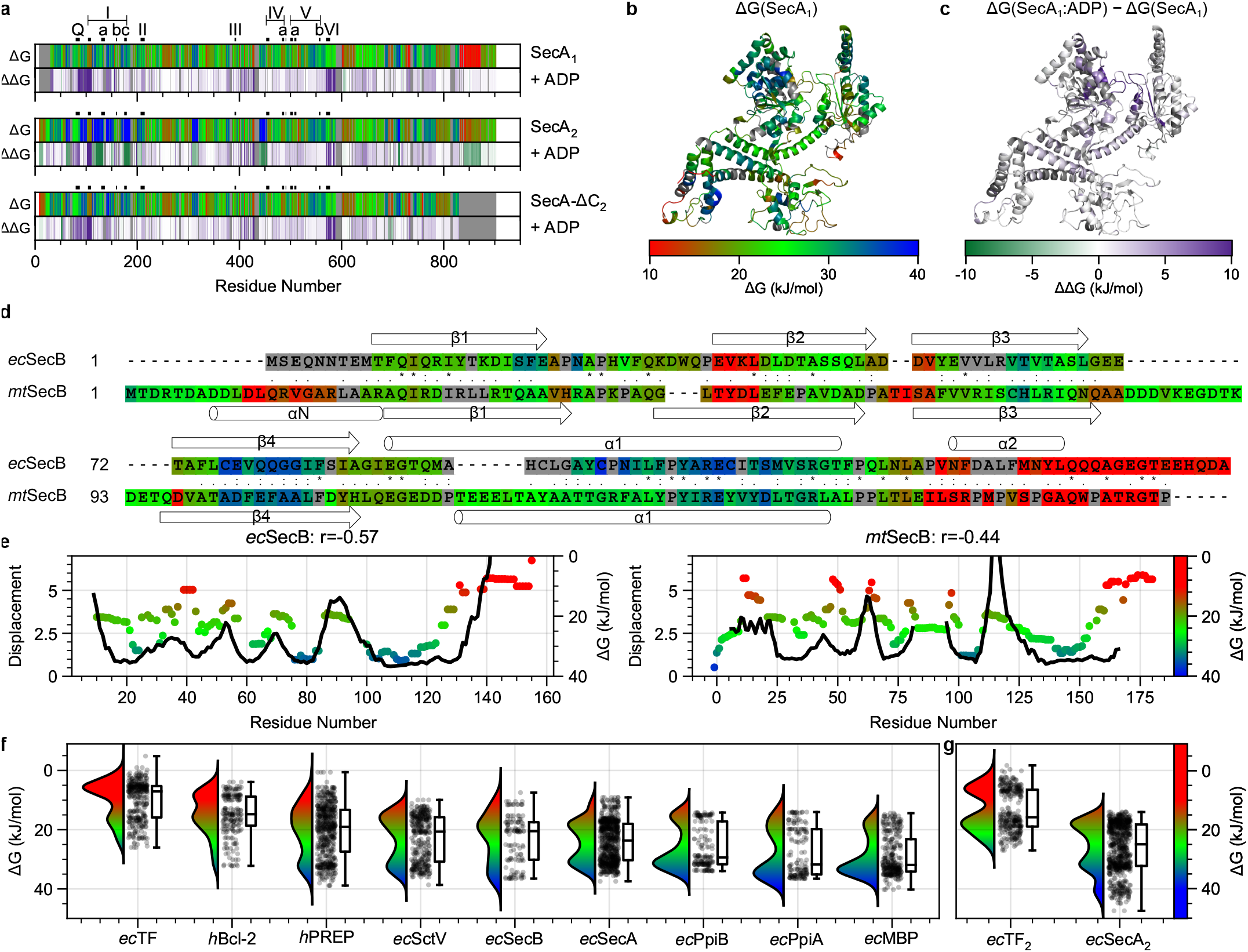
PyHDX analysis applied to a wider protein space. **a**, Protein flexibility of monomeric, dimeric and C-tail deleted SecA as coloured linear bars (Δ*G*, top bars; blue: rigid/red:flexible) and their differential dynamics upon ADP binding (ΔΔ*G*, bottom bars; ADP-rigidified regions in purple). Helicase motifs (Q, I, Ia, Ib, Ic, II, III, IV, IVa, V, Va, Vb and VI; critical regions in ATP hydrolysis and function)[25, 26] are indicated. **b**, Gibbs free energies (Δ*G*) of protein flexibility from HDX-MS for SecA_1_ apoprotein mapped onto its 3D structure (PDB ID 2VDA, ligand removed[27]). **c**, ADP-driven differential dynamics in monomeric SecA shown as differences in Δ*G* (ΔΔ*G*). Regions in purple rigidify upon ADP binding. **d**, Alignment of *ec*SecB (top) and *mt*SecB (bottom) based on both sequence alignment and secondary structure as performed in[28]. Residue similarity is indicated in the middle: * = identical residues,: = strongly similar,. = weakly similar (according to the Gonnet PAM 250 matrix). Colours indicate Δ*G*/residue. **e**, Total displacement of normal modes 7 – 13 (unweighted sum) plotted together with HDX-MS Gibbs free energy (Δ*G*) difference between ‘open’ and ‘closed’ states for *ec*SecB and *mt*SecB. r-values are Pearson correlation coefficients. **f** and **g**, Raincloud plots[29] of Δ*G*/residue for the indicated proteins (**f**) and two dimeric derivatives (**g**).

We next compared two structural homologues, analysed at different experimental conditions: *ec*SecB and *mt*SecB, from *M. tuberculosis* showing modest sequence conservation (13% identity/27% similarity), measured at 20 °C, pD 6, 4 timepoints 10 s – 30 min[28]. Secondary structurebased alignment[28] identified which differences between residue pairs should be minimized by *λ*_2_. Initial apparent differences in the two flexibility profiles (Supplementary Fig. 5a) disappeared after applying a small value of *λ*_2_ = 0.5, (Figure 2d). A Clustal sequence alignment yielded less similar flexibility profiles (Supplementary Fig. 5b). We ensured that *λ*_2_ did not artefactually remove differences, by confirming that flexibility profiles remain different in nonaligned sequences (Supplementary Fig. 5c). These observations imply that protein flexibility might be evolutionarily selected prior to individual amino acids.

We next wondered if our findings relate to protein normal mode analysis (NMA), which are simplified, yet powerful mathematical descriptions of vibrational movements in proteins[30, 31]. Each normal mode describes a simple harmonic oscillation of all atoms in a protein with a characteristic frequency. Linear combinations of normal modes are used in the context of protein flexibility and conformational change[32–34]. Like the Δ*G* profiles (Fig. 2d), the lowest-frequency normal modes of *ec*SecB and *mt*SecB were also conserved (Supplementary Fig. 7), common in superfamilies[35–37]. Δ*G* strongly anticorrelates with normal mode displacement, here derived from the sum of the first six non-trivial normal modes (Pearson’s *r* = −0.57 and *r* = −0.44 for *ec*SecB and *mt*SecB respectively, Fig. 2e). We obtained similar results with other structural twins (not shown). Despite how differently the Linderstrøm-Lang model and normal modes describe protein motions, the agreement in their outputs is remarkable. Presumably, since the Hookean potentials between Cα-atoms within a certain distance cut-off applied in NMA predominate in buried or dense regions, low normal mode displacement coincides with slow D-exchange/high Δ*G* values. This orthogonal analysis may identify specific higher-frequency or sparsely populated, but functionally critical, normal modes. To test the applicability of PyHDX to a wider protein space, we derived Δ*G*/residue values for a total of 9 proteins (see Supplementary Information) and visualized their collective distribution independently of sequence position (Fig. 2f). Δ*G* values of residues indicate a wide range of flexibility (0-40 kJ mol^-1^) being distributed within each protein in two, rarely three, distinct populations, predominantly a rigid and a hyper-flexible one (Fig. 2f), that quantitatively shift between them by additional interactions (e.g. dimerization; Fig. 2g). Protein mean Δ*G* values correlate well to flexibility derived from NMA eigenvalues (*r* = −0.77, Supplementary Fig. 8) and per-residue Δ*G* values are anticorrelated to normal mode displacement for most proteins (Supplementary Table 1). In the primary sequence, rigid and hyper-flexible residues are distributed non-randomly, forming alternating islands, displaying in some cases strong periodicity (e.g. MPB, PpiA and PpiB) revealed by power spectral density analysis (Supplementary Fig. 9). Such non-random distribution could underly protein folding and allostery. In summary, PyHDX rapidly processes single and multiple HDX-MS datasets and visualizes residue-level Gibbs free energies on linear and 3D structures. Residue level energies together with normal modes open up previously unavailable possibilities for evolutionary, structural and functional studies and a universal description of protein flexibility.

## Methods

### Protein Purification

*ec*SecB_4_ was purified as previously described[38]. *ec*SecB_2_ was generated by mutating 3 residues (Y109A, T115A, S119A) that form the dimer-dimer interface and purified by nickel affinity purification as described[38]. Mutations were introduced by Quick-Change Mutagenesis System (Pfu turbo, Agilent) using plasmid pIMBB490 (pET16b secB) as template. Mutations were introduced using the primers in Table 1.

**Table 1:**
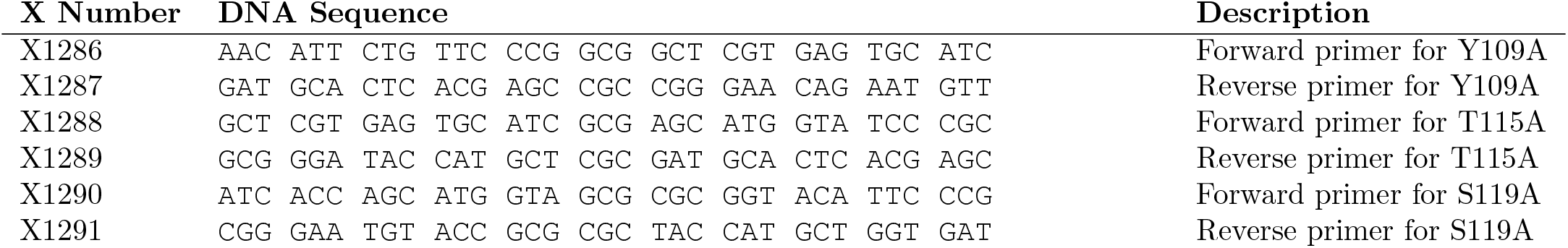
Primers for generating the *ec*SecB_2_ mutant

### Hydrogen/deuterium exchange mass spectrometry

D-exchange was initiated by diluting 100 pmol of SecB by 10 fold into D_2_O buffer (50 mM Tris-HCl pH_read_ 8.0, 50 mM KCl, 1 mM MgCl_2_, 4 μM ZnSO_2_, 2 mM TCEP) reconstituted in 99.9% D_2_O (Euroiso-top) to obtain a final D_2_O concentration of 90%. Continuous deuterium labelling was carried out for 6 timepoints (10 s, 30 s, 1 min, 5 min, 10 min, 100 min) at 30 °C. The reaction was quenched by the addition of pre-chilled quench buffer (1.5% formic acid, 4 mM TCEP, 0.1% DDM) in a 1:1 ratio. Reaction was injected into a nanoAcquity UPLC system with HDX technology (Waters, UK) coupled to a Synapt G2 ESI-QTOF mass spectrometer. Protein digestion was carried out by an online home-packed immobilized pepsin (Sigma) column (2mm x 2 cm, Idex) at 18 °C. LC and instrument parameters were as previously described[39]. The 100% deuteration control (FD control) was obtained by incubating SecB in D_2_O buffer containing 6 M UreaD_4_ (98% D, Sigma) overnight at room temperature. Peptide identification was carried out in ProteinLynx Global Server (PLGS, Waters UK) and deuterium exchange data was analysed in DynamX 3.0 (Waters, UK). ‘State data’ was exported from DynamX in the form of csv files. *mt*SecB HDX-MS data was corrected for back-exchange by assuming a constant 28% back-exchange for all peptides. HDX-MS data of other proteins were from previously published datasets (hPREP[39], *ec*SecA[25]) or will be described in detail elsewhere (*ec*TF, *h*Bcl-2, *ec*SctV *ec*PpiB, *ec*PpiA and *ec*MBP).

### Theory

The Linderstrøm-Lang model describing HDX-MS experiments can be written as[8, 9]:

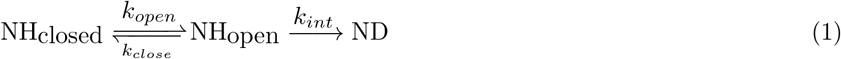

Where NH represents an amide hydrogen and ND an amide deuterium. The intrinsic exchange rate *k_int_* (frequently also referred to as chemical exchange rate, *k_ch_*) dependent on the pH and temperature at which the deuterium labelling reaction takes place, as well as the primary sequence of the peptide, and can be calculated accurately[40–43]. This intrinsic rate is a major influence on the observed kinetics of D-exchange as it can vary up to three orders of magnitude (e.g. pH 6, 0°C, vs pH 8, 30°C). To correct for back-exchange[3, 44], a fully deuterated (FD) control sample is used. The experimentally determined FD provides the maximal degree of D-exchange possible for any given peptide. The corrected D-uptake for each peptide is then calculated as[44]:

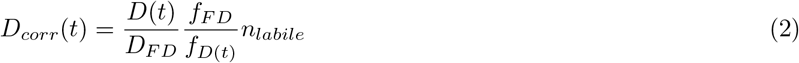

Where *D* is the experimentally measured D-uptake, *D_FD_* the D-uptake of the fully deuterated control and *f_FD_* and *f_FD_*(*t*) represent the fractional D-content of the FD control buffer and the D-labelling buffer, respectively. *n_labile_* is the number of exchange-competent amide groups in the peptide. Using the steady-state approximation, the observed rate of formation of the deuterated residue ND is given by[45]:

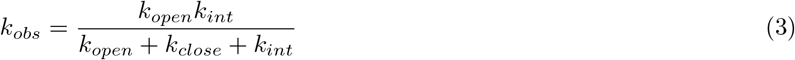

Assuming that the protein dynamics are faster than the exchange reaction (*k_open_* + *k_close_* ≫ *k_int_*) and introducing the substitution *PF* = *k_close_/k_open_*, the expression reduces to:

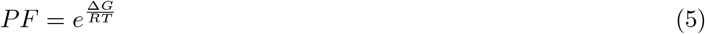

Where *PF* is the protection factor[9, 20] for this particular protein residue. This ratio of rates is equivalent to the system’s Boltzmann factor and thus relates to the Gibbs free energy difference between the closed and open states:

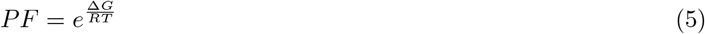

Where the sign of Δ*G* is chosen such that Δ*G* is positive when the energy of the open state is higher compared to the closed state, as is generally true for structured proteins. A low value of Δ*G* indicates highly dynamic or disordered proteins. We can now write down the Lagrangian (cost function) for finding Δ*G*. Given a protein with *N_r_* residues *r* and a HDX-MS experiment which yielded *N_p_* peptides *p* at *N_t_* timepoints *t*, each with an associated measured deuterium uptake *D*(*t, p*) (Table 2), the Lagrangian is:

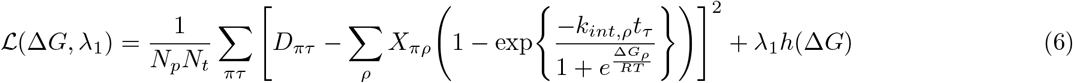

**Table 2:** Lagrangian symbols and indices

| Axis | Number of elements | Index |
| --- | --- | --- |
| Samples | $N_s$ | $\sigma$ |
| Peptides | $N_p$ | $\pi$ |
| Residues | $N_r$ | $\rho$ |
| Timepoints | $N_t$ | $\tau$ |

With:

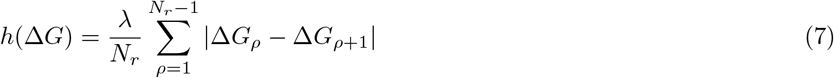

Where *D* is the corrected D-uptake and *X* is a ‘coupling matrix’ describing to which residues each peptide corresponds:

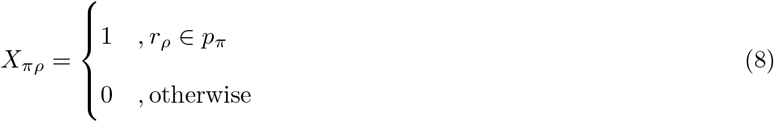

Such that its elements are 1 when the corresponding residues are found in a given peptide. When expanding to global fitting of *N_s_* HDX-MS samples (e.g. liganded, oligomers, homologues, pH/temperature variation) the Lagrangian becomes:

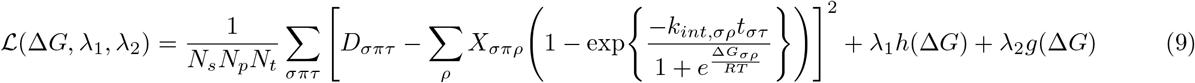

With

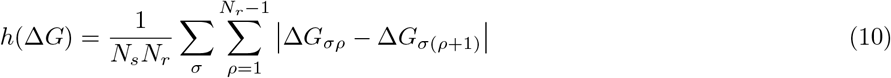

And

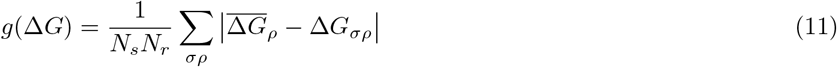

Covariances are obtained from the diagonal elements of the inverse of the Hessian:

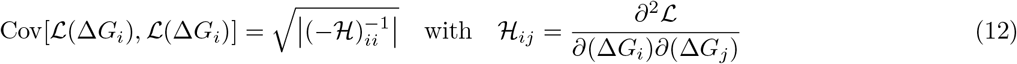

### Implementation

In PyHDX, all quantities are PyTorch[45] Tensors or Numpy[46] arrays, with shapes as indicated in Table 3, such that the Lagrangian can be computed through matrix multiplications, where in the 3D case multiplication is done in batch along the first axis, according to Python’s PEP465 convention. For minimization of the Lagrangian, we use the PyTorch machine learning framework, such that its autograd automatic differentiation engine can be used to accelerate the process. The Stochastic-Gradient Descent (SGD) method is used by default (learning rate: 10, momentum: 0.5, nesterov: True) To further ensure convergence to the correct solution, the Δ*G* is initialized with guess values obtained from weighted averaging (by inverse peptide length) of all peptides for a given timepoint. This procedure yields a kinetic uptake curve per residue from which apparent exchange rates are determined.

**Table 3:** Lagrangian variables and their shapes

| Symbol | Description | Shape | Batch shape |
| --- | --- | --- | --- |
| $D$ | D-uptake values | $N_p \times N_t$ | $N_s \times N_p \times N_t$ |
| $X$ | Coupling matrix | $N_p \times N_r$ | $N_s \times N_p \times N_r$ |
| $k_{int}$ | Intrinsic exchange rates | $N_r \times 1$ | $N_s \times N_r \times 1$ |
| $\Delta G$ | Gibbs free energy | $N_r \times 1$ | $N_s \times N_r \times 1$ |
| $t$ | Timepoints | $1 \times N_r$ | $N_s \times 1 \times N_t$ |
| $T$ | Temperature | $1 \times 1$ | $N_s \times 1 \times 1$ |

PyHDX was built on top of the scientific python ecosystem. Computation is done using the packages numpy[46], pandas[47], scipy[48], scikit-image[49] and symfit[50]. Fitting of Δ*G* is implemented on the machine learning platform PyTorch[51]. Computationally intensive tasks are scheduled to be processed in parallel through Dask[52]. Intrinsic exchange rates are calculated as previously described[40–40] and implemented by HDXRate[53]. Graphical output is generated with either matplotlib[54] or bokeh[55]. PyHDX features an API for data analysis in Jupyter notebooks[56] and a web application implemented in panel[57] using NGL[58, 59] to visualize proteins.

### Power Spectral Density

To calculate PSDs, we first filled missing values in the Δ*G* linear map by interpolation. Next, the DC component was removed by subtracting mean Δ*G*. The resulting signal was autocorrelated, normalized and Fourier transformed. Periodicity was determined from the peak in the resulting PSD.

### Normal mode analysis

Normal modes are calculated using the WebNM@ web server[60]. Input structural models used are shown in Supplementary Table 1. For multi-chain oligomeric proteins, per-residue displacements are averaged across protein chains. Displacements per normal mode are then summed with equal weights for the first 6 non-trivial normal modes. Normal mode flexibility is derived from normal mode eigenvalues as described previously[32]:

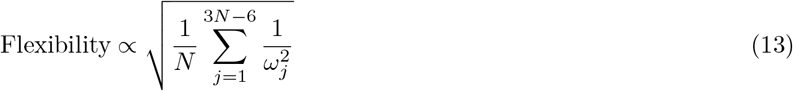

Where *N* is the number of residues and *ω_j_* is the frequency of normal mode *j*.

## Data Availability

*ec*SecB HDX-MS state data is available on the PyHDX GitHub repository. SecA HDX-MS state data was published previously[25].

## Acknowledgements

We are grateful to: J. Marcoux, P. Geneveux and L. Mourey for generously sharing *mt*SecB HDX-MS data; J. Claesen for discussions; A. Portaliou for constructs; the open source software community for their support and advice. Research in our lab was funded by grants (to AE): ProFlow (FWO/F.R.S.-FNRS “Excellence of Science - EOS” programme grant #30550343), CARBS (#G0C6814N; FWO) and PROFOUND (WOG; W002421N; FWO) and (to AE and SK): FOscil (ZKD4582 - C16/18/008; KU Leuven). SKr was a FWO [PEGASUS]_2_ MSC postdoctoral fellow. JHS is a PDM, KU Leuven postdoctoral fellow. This project has received funding from the Research Foundation – Flanders (FWO) and the European Union’s Horizon 2020 research and innovation programme under the Marie Skłodowska-Curie grant agreement No 665501.

## Author Contributions

JHS conceived all mathematical analysis and developed and implemented software and web interface. SKr, BYS, RP and SK provided HDX-MS data and analysis. SKr guided optimization of software parameters and validated output. JHS wrote the first draft with contributions from SKr and AE. All authors reviewed and approved the final manuscript. AE conceived and managed the project.

## Supplementary Information

**Supplementary Figure 1:**
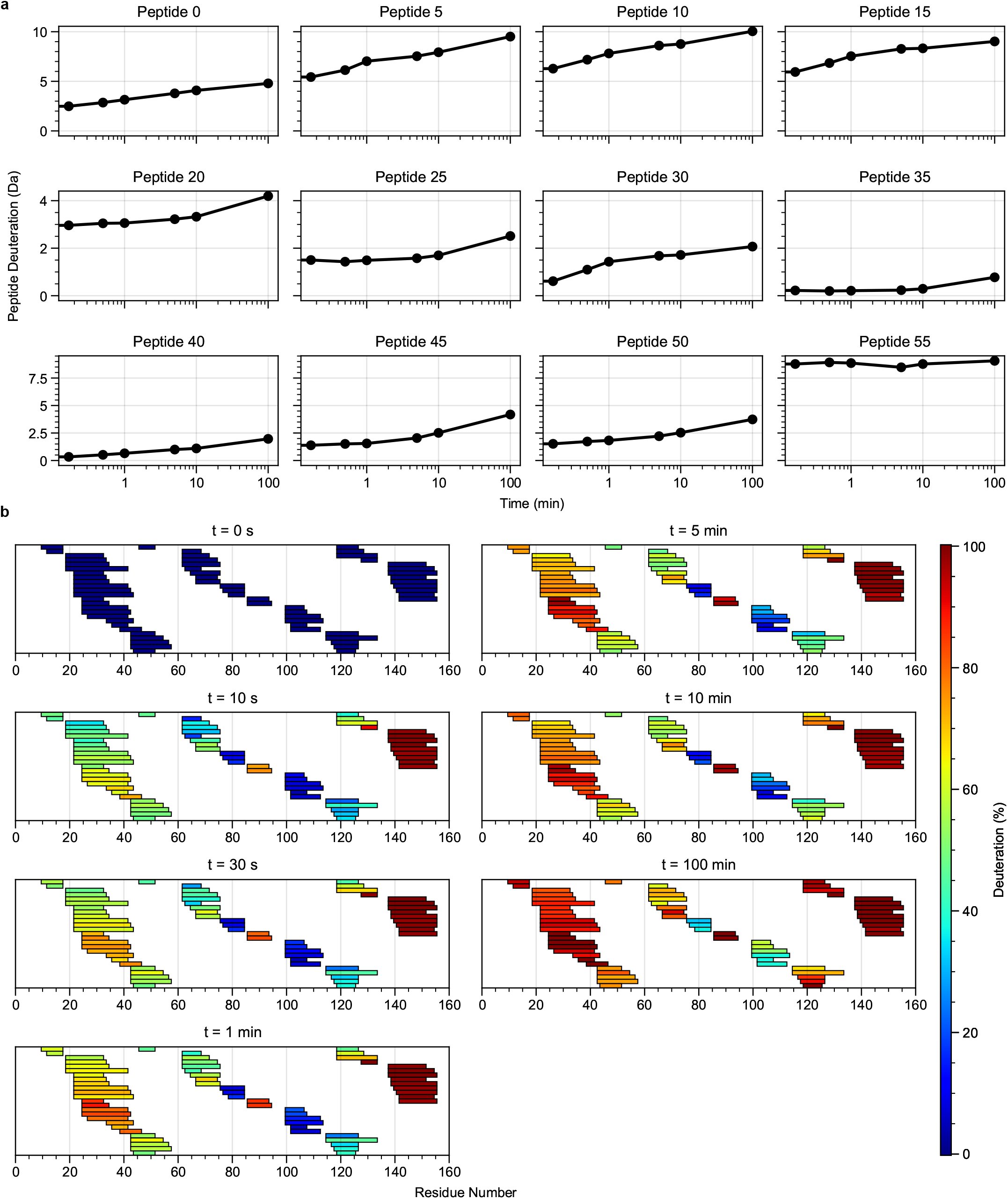
Example of HDX-MS peptide input data. **a**, Uncorrected deuterium uptake plots for selected peptides obtained for *ec*SecB. **b**, Peptide heatmaps for all timepoints showing relative deuterium uptake with respect to a fully deuterated (FD) control sample.

**Supplementary Figure 2:**
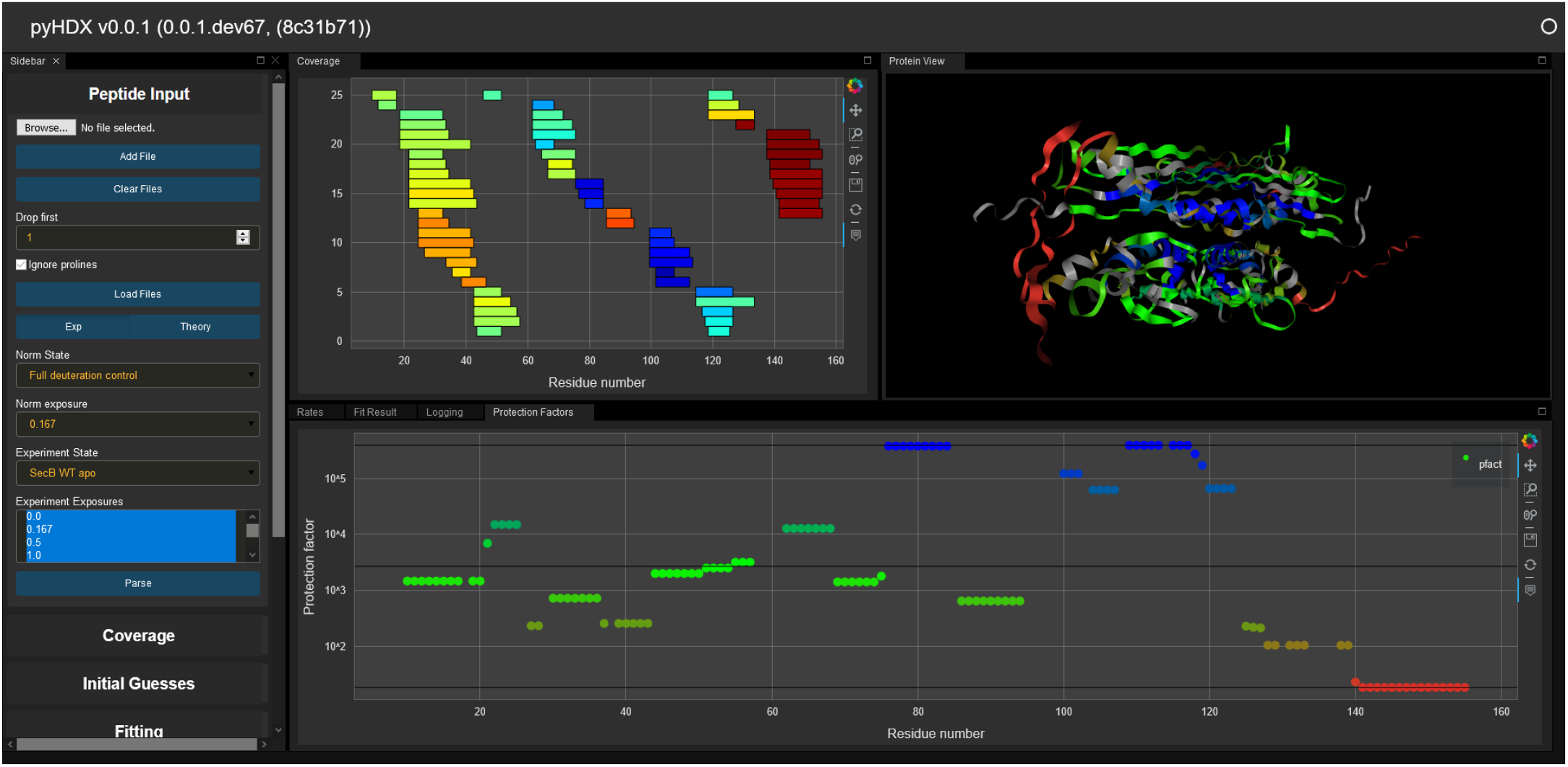
Screenshot of PyHDX web interface. The web interface is publicly accessible and researchers can use it to upload data, apply FD control corrections, determine Δ*G* initial guesses, determine Δ*G* by global fit of all peptides. Results are directly visualized in the web interface either as linear map or coloured 3D structure, or can be exported in .csv format and pymol-colouring scripts.

**Supplementary Figure 3:**
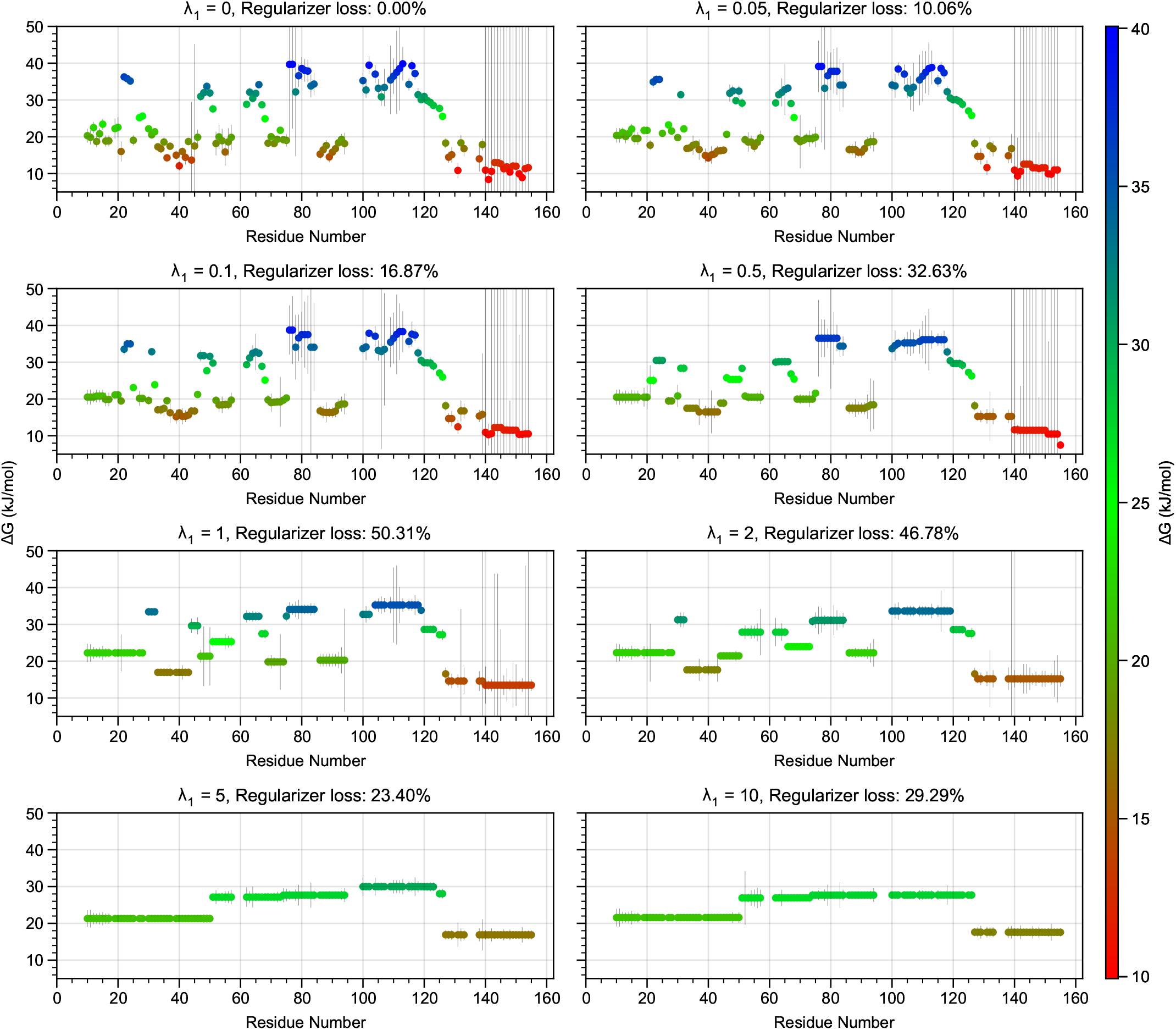
Influence of regularizer *λ*_1_ on the obtained Δ*G* values for *ec*SecB_4_. At *λ*_1_ = 0 Δ*G* can vary without penalty along the residue axis resulting in scattered Δ*G*. Regularizer loss percentages are fraction of total Lagrangian value due to regularization terms. Recommended values for regularization percentages are around 10%. At too high values for *λ*_1_ the figure shows that features in Δ*G* are dampened out. Error bars are covariances derived from the Hessian of the unconstraint Lagrangian.

**Supplementary Figure 4:**
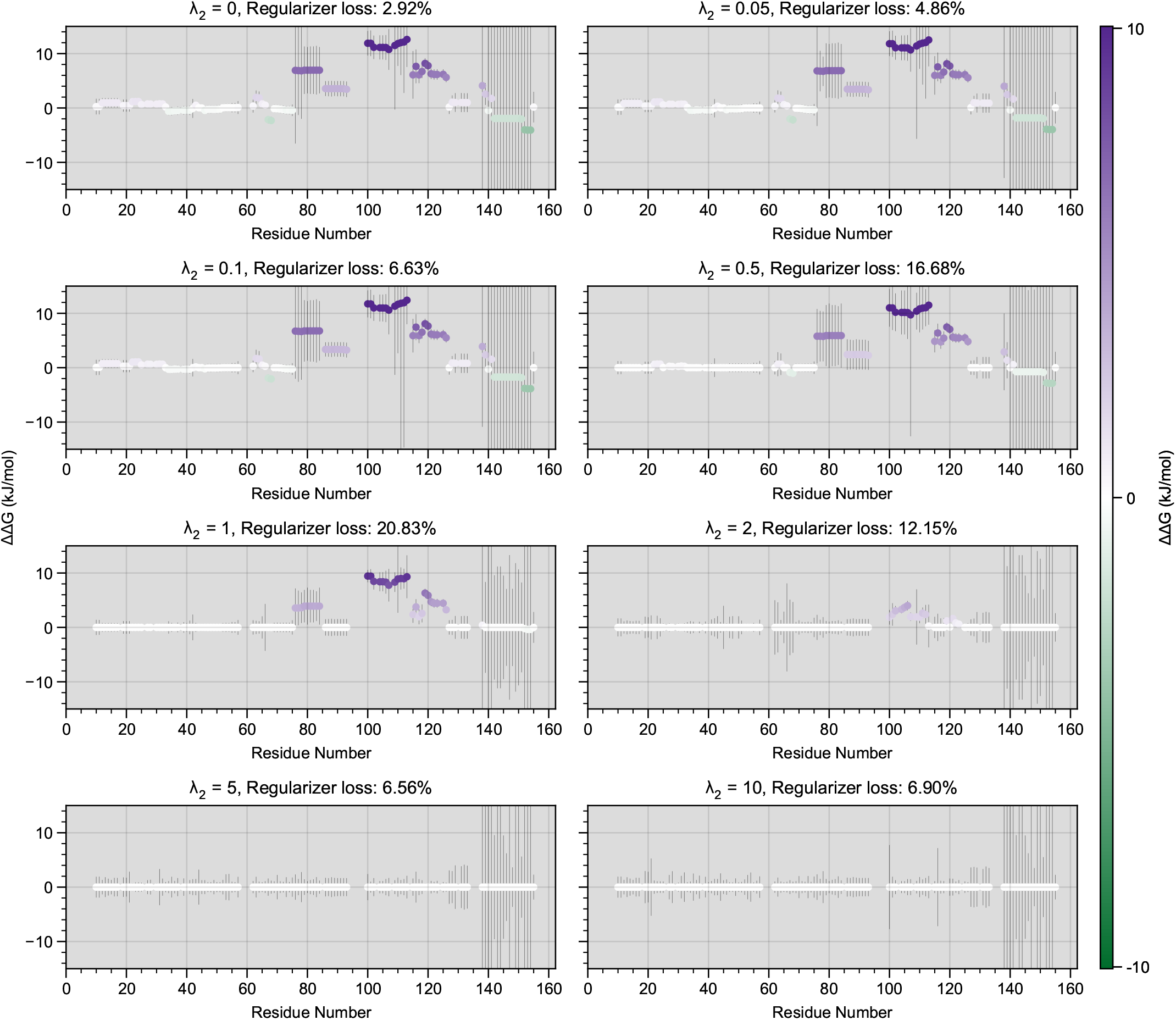
Influence of regularizer *λ*_2_ on the obtained ΔΔ*G* values between *ec*SecB_4_ and *ec*SecB_2_. Purple colour and positive ΔΔ*G* values represents local rigidity of *ec*SecB_4_ compared to *ec*SecB_2_, negative ΔΔ*G* values and green colour represents local flexibility. As *λ*_2_ constraints differences between datasets, low values will show maximum differences, potentially leading to artefactual “differences” between datasets, while high values of *λ*_2_ select only the most significant differences between datasets. Error bars are propagated covariances from *ec*SecB_4_ and *ec*SecB_2_. Regularizer loss percentages are fraction of total Lagrangian value due to regularization terms.

**Supplementary Figure 5:**
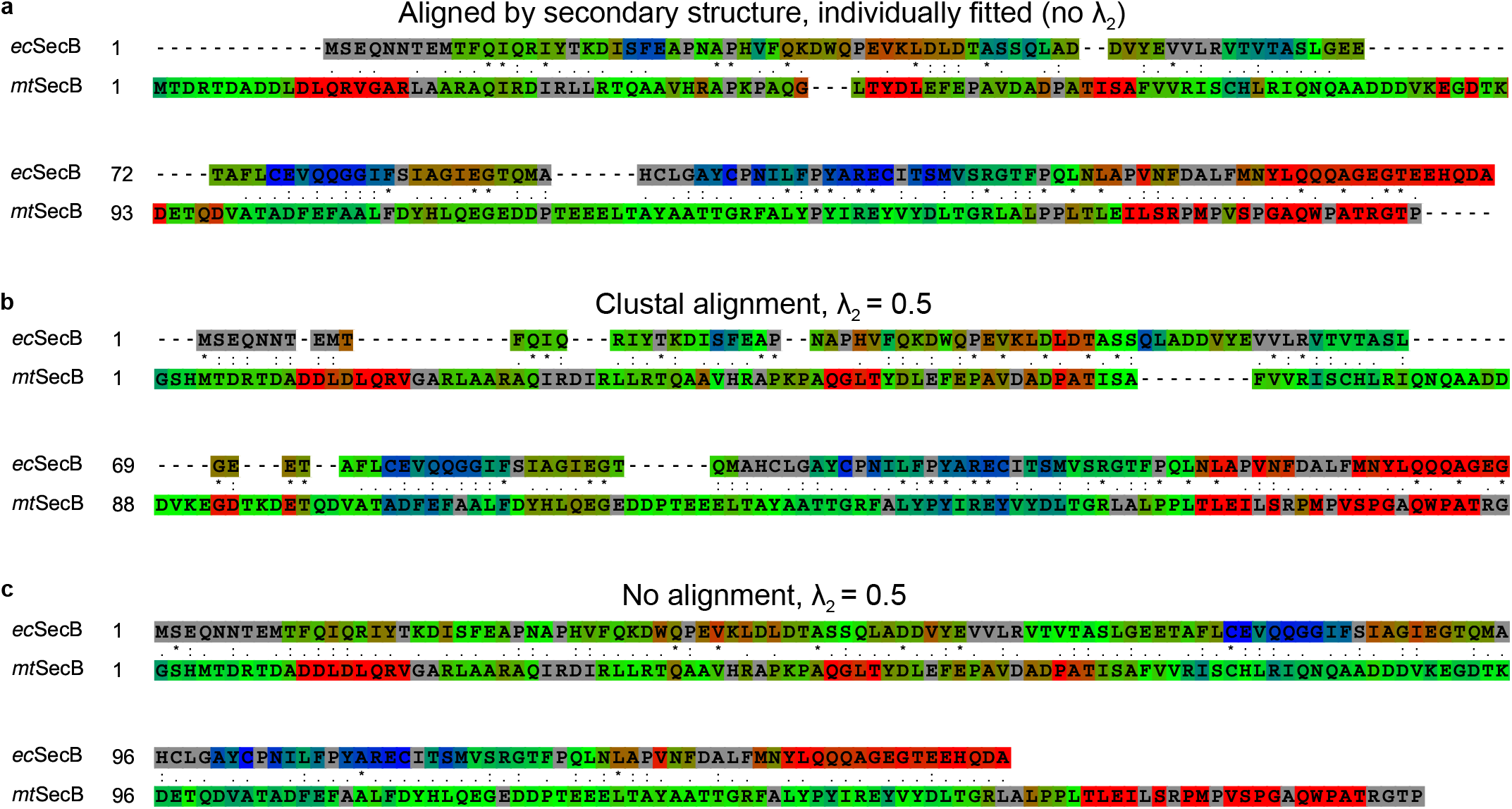
Comparison of *ec*SecB and *mt*SecB Δ*G* with alternative aligned fitting strategies. Regularizer *λ*_1_ = 0.05 for all panels. Residue similarity is indicated in the middle: * = identical residues,: = strongly similar,. = weakly similar (according to the Gonnet PAM 250 matrix). **a**, Proteins are aligned by secondary structure[1], fitting is done individually for both proteins (no *λ*_1_). **b**, No alignment (aligned by residue number), *λ*_2_ = 0.5 **c**, Proteins were aligned by Clustal Omega (ClustalW with character count)[2], *λ*_2_ = 0.5.

**Supplementary Figure 6:**
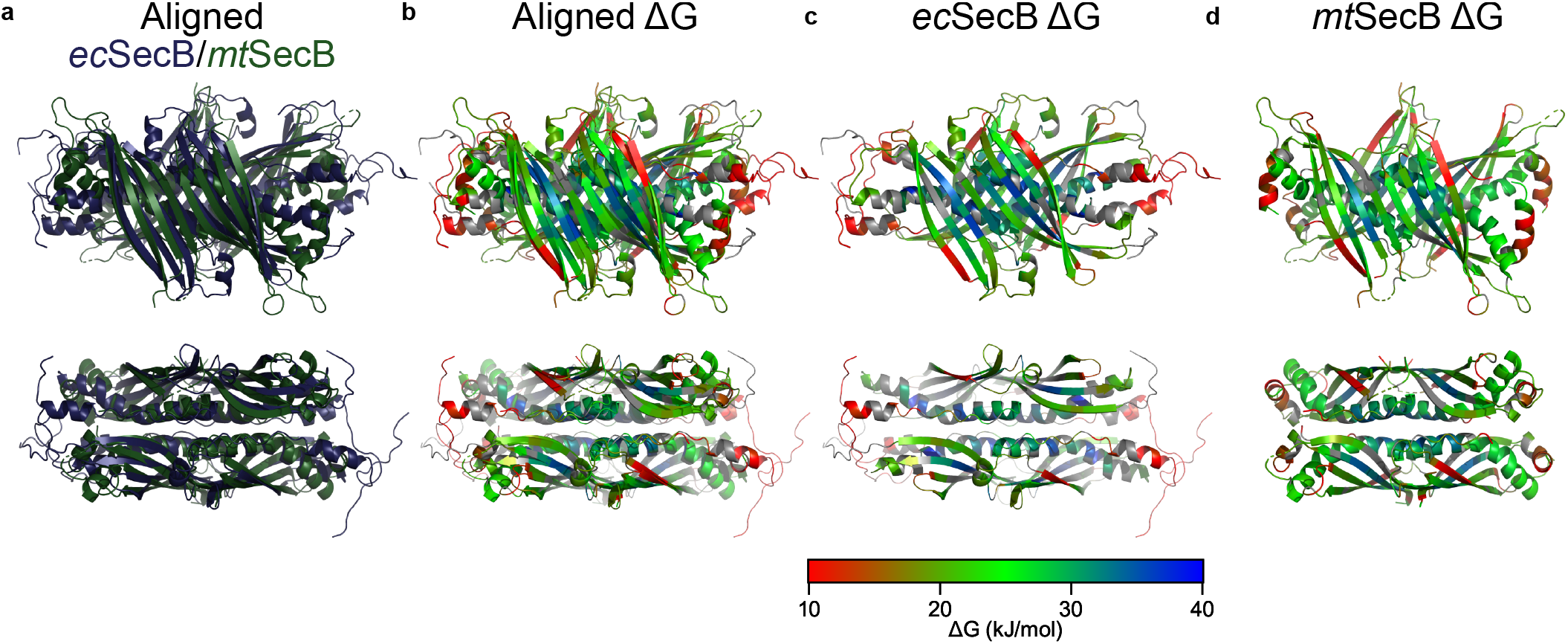
Structural alignment of homologous proteins *ec*SecB and *mt*SecB by tertiary structure. (Pymol “align” command) **a**, Aligned structures coloured by protein (*ec*SecB; Blue, *mt*SecB; Green). **b**, Aligned structures coloured by Δ*G*, showing a high degree of flexibility conservation in 3D space between the two proteins. **c**, **d**, Structures of individual proteins in identical orientation compared to aligned structures, coloured by Δ*G*.

**Supplementary Figure 7:**
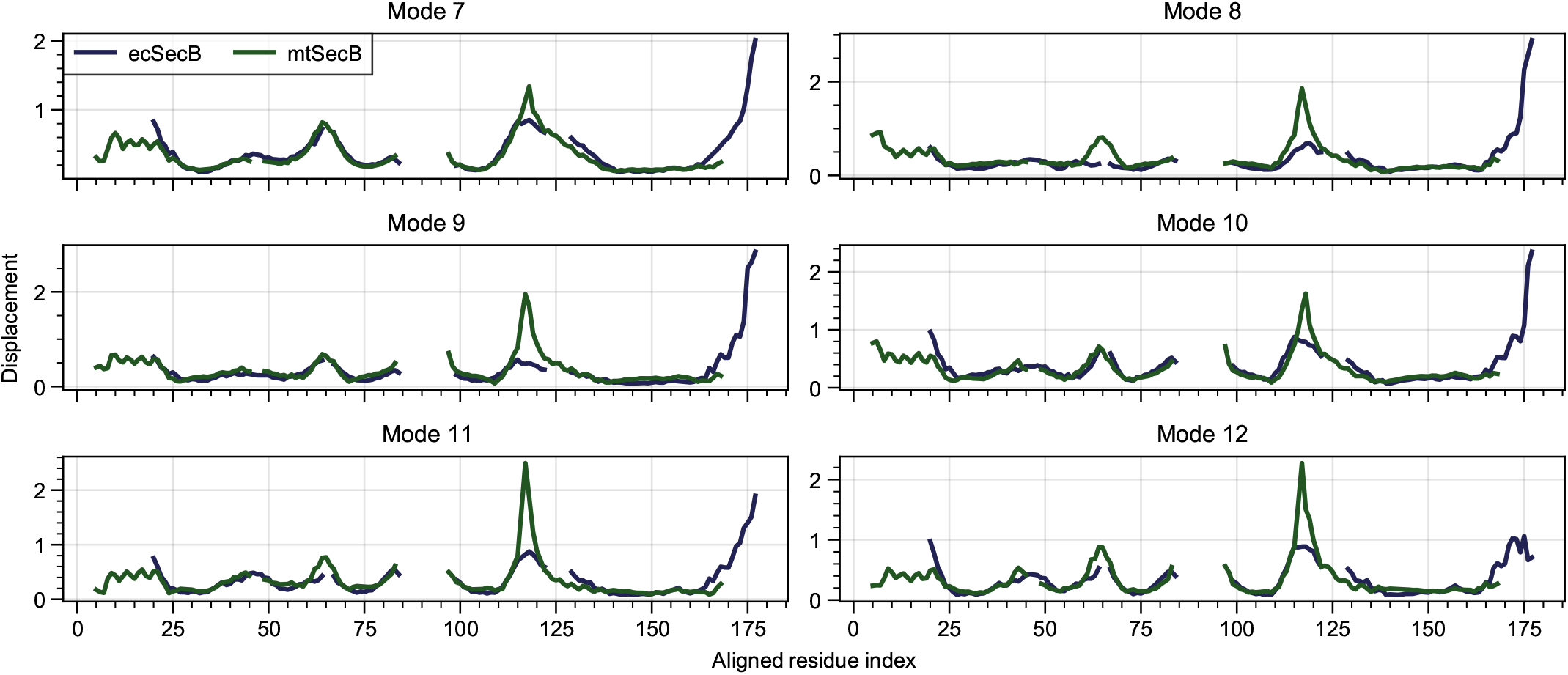
Alignment of *ec*SecB and *mt*SecB normal modes[3]. The figure shows Cα displacements for the first 6 non-trivial normal modes. Residues are aligned by secondary structure as described in[1]

**Supplementary Figure 8:**
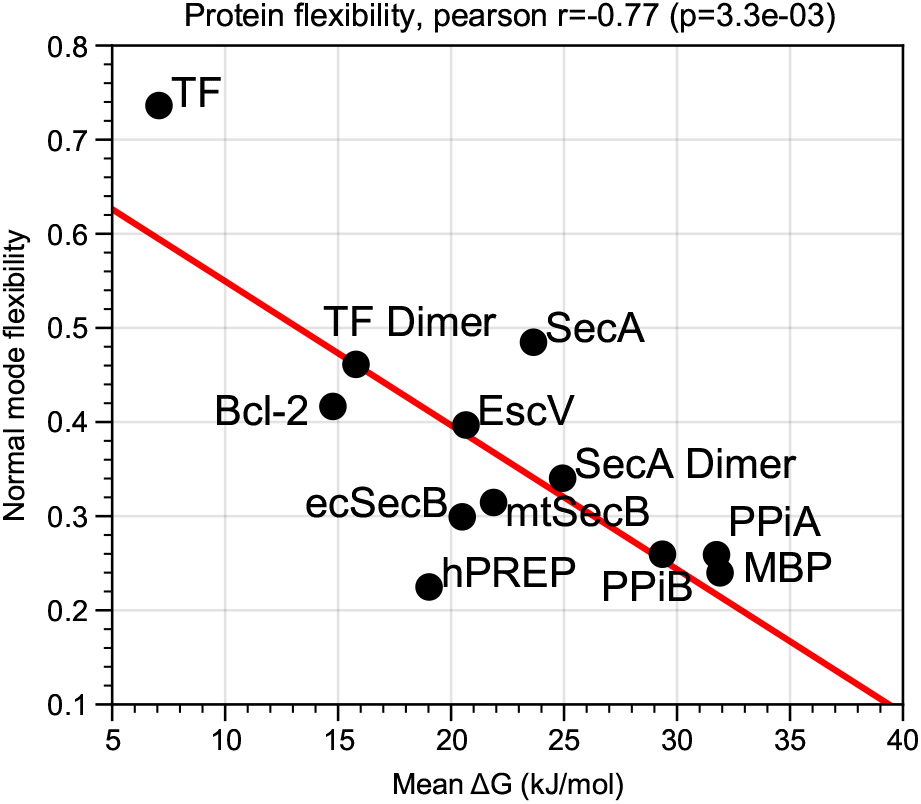
Flexibility derived from normal mode analysis versus mean Δ*G* of selected proteins. Normal mode flexibility is calculated from the eigenvalues of the first 200 normal modes[4] as described in the Methods section.

**Supplementary Figure 9:**
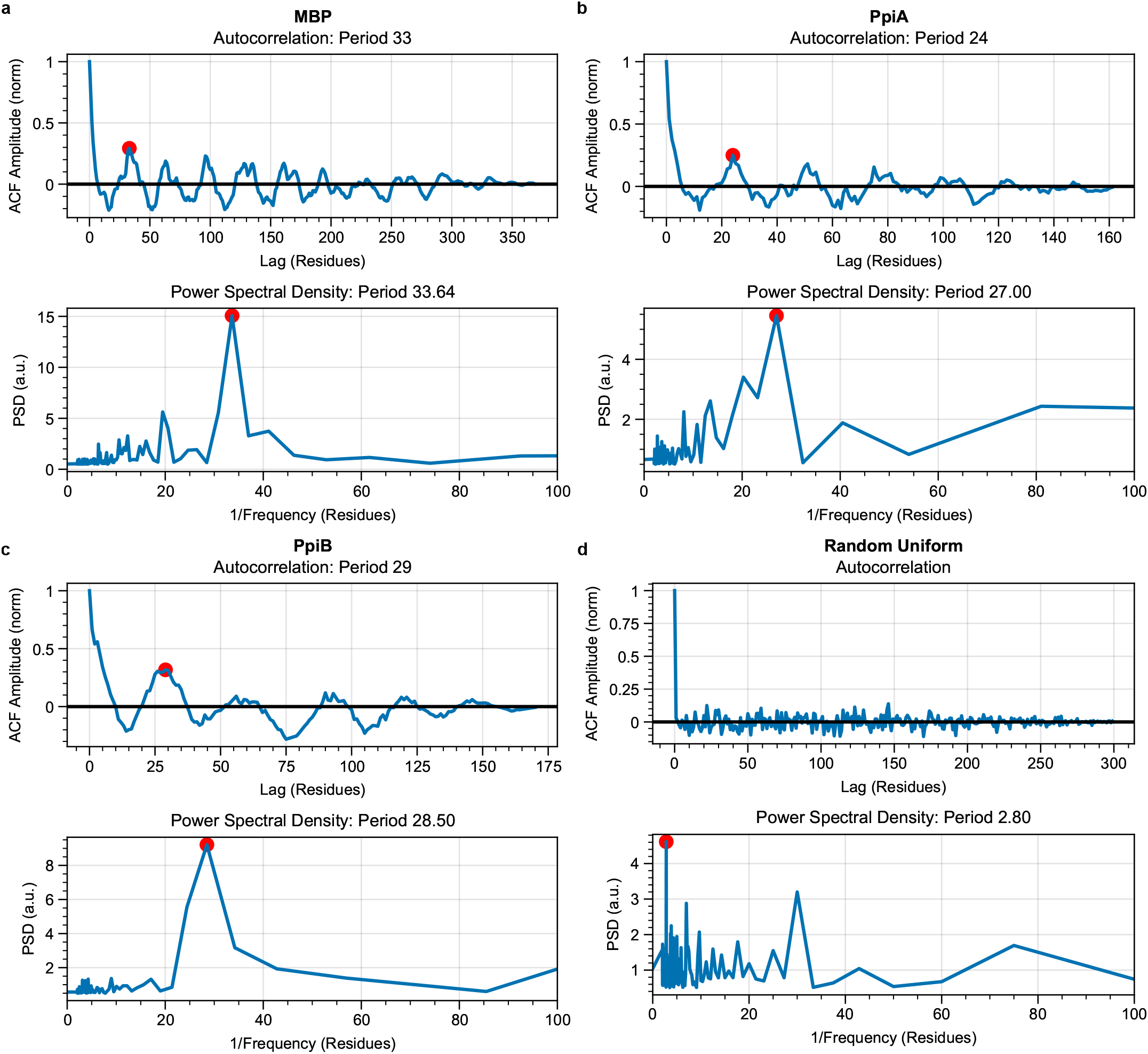
Power spectral density (PSD) analysis of periodicity in Δ*G* along primary sequence. Top panels show normalized autocorrelated signal (ACF) where the position of the first peak is highlighted. Bottom panels show Fourier transform of ACF (=PSD) plotted as a function of inverse frequency. Panels shown are **a**, *ec*MBP. **b**, *ec*PpiA, **c**, *ec*PpiB, **d**, Control PSD analysis on 300 random data points uniformly sampled from the [0, 1) interval, showing no long-range correlations.

**Supplementary Table 1:**
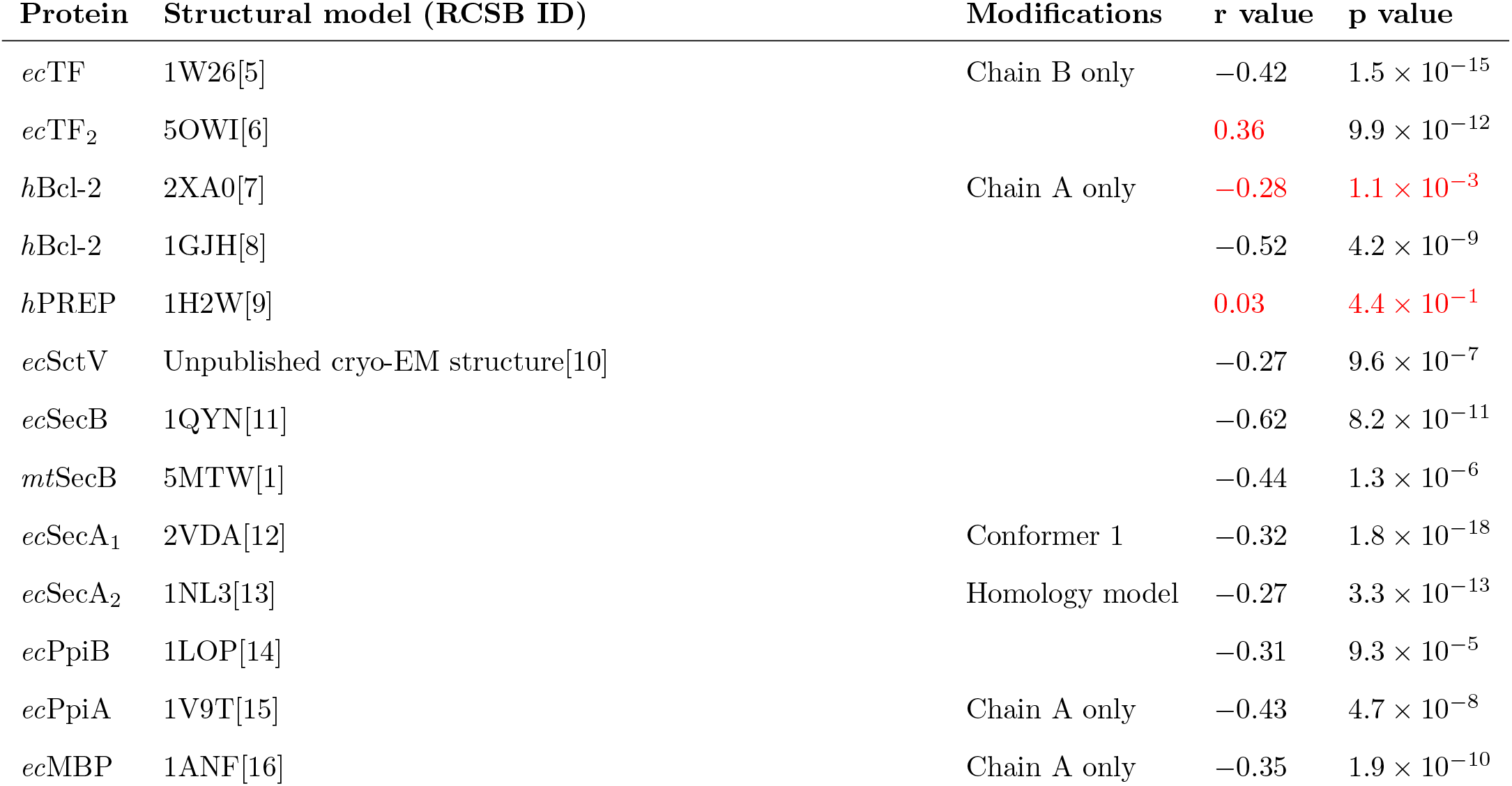
Correlations between Δ*G* values and displacement in the 6 lowest frequency normal modes of selected proteins. For all structural models, any ligand is removed from the structure when applicable. The sign of the correlation value is expected to be negative if Δ*G* correlates with normal mode displacement as high Δ*G* values correspond to rigid (low displacement) regions and low Δ*G* values correspond to flexible (high displacement) regions. Poorly anticorrelating entries are highlighted in red.

| Protein | Structural model (RCSB ID) | Modifications | r value | p value |
| --- | --- | --- | --- | --- |
| *ec*TF | 1W26[5] | Chain B only | -0.42 | $1.5 \times 10^{-15}$ |
| *ec*TF<sub>2</sub> | 5OWI[6] | | **0.36** | $9.9 \times 10^{-12}$ |
| *h*Bcl-2 | 2XA0[7] | Chain A only | **-0.28** | **$1.1 \times 10^{-3}$** |
| *h*Bcl-2 | 1GJH[8] | | -0.52 | $4.2 \times 10^{-9}$ |
| *h*PREP | 1H2W[9] |  | **0.03** | **$4.4 \times 10^{-1}$** |
| *ec*SctV | Unpublished cryo-EM structure[10] | | -0.27 | $9.6 \times 10^{-7}$ |
| *ec*SecB | 1QYN[11] | | -0.62 | $8.2 \times 10^{-11}$ |
| *mt*SecB | 5MTW[1] | | -0.44 | $1.3 \times 10^{-6}$ |
| *ec*SecA<sub>1</sub> | 2VDA[12] | Conformer 1 | -0.32 | $1.8 \times 10^{-18}$ |
| *ec*SecA<sub>2</sub> | 1NL3[13] | Homology model | -0.27 | $3.3 \times 10^{-13}$ |
| *ec*PpiB | 1LOP[14] | | -0.31 | $9.3 \times 10^{-5}$ |
| *ec*PpiA | 1V9T[15] | Chain A only | -0.43 | $4.7 \times 10^{-8}$ |
| *ec*MBP | 1ANF[16] | Chain A only | -0.35 | $1.9 \times 10^{-10}$ |

## References

[1] Karplus, M. & Kuriyan, J. Molecular dynamics and protein function. Proceedings of the National Academy of Sciences of the United States of America 102, 6679–6685 (2005).

[2] Fajer, P. G., Bou-Assaf, G. M. & Marshall, A. G. Improved Sequence Resolution by Global Analysis of Overlapped Peptides in Hydrogen/Deuterium Exchange Mass Spectrometry. Journal of The American Society for Mass Spectrometry 23, 1202–1208 (2012).

[3] Masson, G. R. et al. Recommendations for performing, interpreting and reporting hydrogen deuterium exchange mass spectrometry (HDX-MS) experiments. Nature Methods 16, 595–602 (2019).

[4] Oganesyan, I., Lento, C. & Wilson, D. J. Contemporary hydrogen deuterium exchange mass spectrometry. Methods 144, 27–42 (2018).

[5] Walters, B. T., Ricciuti, A., Mayne, L. & Englander, S. W. Minimizing Back Exchange in the Hydrogen Exchange-Mass Spectrometry Experiment. Journal of the American Society for Mass Spectrometry 23, 2132–2139 (2012).

[6] Claesen, J. & Burzykowski, T. Computational methods and challenges in hydrogen/deuterium exchange mass spectrometry. Mass Spectrometry Reviews 36, 649–667 (2017).

[7] Vadas, O. & Burke, J. Probing the dynamic regulation of peripheral membrane proteins using hydrogen deuterium exchange–MS (HDX–MS). Biochemical Society Transactions 43, 773–786 (2015).

[8] Linderstrøm-Lang, K. Deuterium exchange between peptides and water. Chem. Soc. Spec. Publ. 2, 1–20 (1955).

[9] Englander, S. W., Sosnick, T. R., Englander, J. J. & Mayne, L. Mechanisms and uses of hydrogen exchange. Current Opinion in Structural Biology 6, 18–23 (1996).

[10] Huang, C., Rossi, P., Saio, T. & Kalodimos, C. G. Structural basis for the antifolding activity of a molecular chaperone. Nature 537, 202–206 (2016).

[11] Gessner, C. et al. Computational method allowing Hydrogen-Deuterium Exchange Mass Spectrometry at single amide Resolution. Scientific Reports 7, 1–10 (2017).

[12] Skinner, S. P., Radou, G., Tuma, R., Houwing-Duistermaat, J. J. & Paci, E. Estimating Constraints for Protection Factors from HDX-MS Data. Biophysical Journal 116, 1194–1203 (2019).

[13] Saltzberg, D. J. et al. A Residue-Resolved Bayesian Approach to Quantitative Interpretation of Hydrogen-Deuterium Exchange from Mass Spectrometry: Application to Characterizing Protein–Ligand Interactions. The Journal of Physical Chemistry B 121, 3493–3501 (2017).

[14] Salmas, R. E. & Borysik, A. J. HDXmodeller: an online webserver for high-resolution HDX-MS with auto-validation. Communications Biology 4, 1–8 (2021).

[15] Althaus, E. et al. Computing H/D-Exchange rates of single residues from data of proteolytic fragments. BMC Bioinformatics 11, 424 (2010).

[16] Zhang, Z., Zhang, A. & Xiao, G. Improved Protein Hydrogen/Deuterium Exchange Mass Spectrometry Platform with Fully Automated Data Processing. Analytical Chemistry 84, 4942–4949 (2012).

[17] Zhang, Z. Complete Extraction of Protein Dynamics Information in Hydrogen/Deuterium Exchange Mass Spectrometry Data. Analytical Chemistry 92, 6486–6494 (2020).

[18] Hamuro, Y. & Zhang, T. High-Resolution HDX-MS of Cytochrome c Using Pepsin/Fungal Protease Type XIII Mixed Bed Column. Journal of The American Society for Mass Spectrometry 30, 227–234 (2019).

[19] Kan, Z.-y., Ye, X., Skinner, J. J., Mayne, L. & Englander, S. W. ExMS2: An Integrated Solution for Hydrogen–Deuterium Exchange Mass Spectrometry Data Analysis. Analytical Chemistry 91, 7474–7481 (2019).

[20] Roder, H., Elöve, G. A. & Englander, S. W. Structural characterization of folding intermediates in cytochrome c by H-exchange labelling and proton NMR. Nature 335, 700–704 (1988).

[21] Englander, S. W. & Kallenbach, N. R. Hydrogen exchange and structural dynamics of proteins and nucleic acids. Quarterly Reviews of Biophysics 16, 521–655 (1983).

[22] Pirrone, G. F., Iacob, R. E. & Engen, J. R. Applications of Hydrogen/Deuterium Exchange MS from 2012 to 2014. Analytical Chemistry 87, 99–118 (2015).

[23] Chalmers, M. J., Busby, S. A., Pascal, B. D., West, G. M. & Griffin, P. R. Differential hydrogen/deuterium exchange mass spectrometry analysis of protein–ligand interactions. Expert Review of Proteomics 8, 43–59 (2011).

[24] Sala, A., Bordes, P. & Genevaux, P. Multitasking SecB chaperones in bacteria. Frontiers in Microbiology 5 (2014).

[25] Krishnamurthy, S. et al. A nexus of intrinsic dynamics underlies translocase priming. Structure (2021).

[26] Papanikou, E., Karamanou, S. & Economou, A. Bacterial protein secretion through the translocase nanomachine. Nature Reviews Microbiology 5, 839–851 (2007).

[27] Gelis, I. et al. Structural Basis for Signal-Sequence Recognition by the Translocase Motor SecA as Determined by NMR. Cell 131, 756–769 (2007).

[28] Guillet, V. et al. Structural insights into chaperone addiction of toxin-antitoxin systems. Nature Communications 10, 782 (2019).

[29] Allen, M. et al. Raincloud plots: a multi-platform tool for robust data visualization. Wellcome Open Research 4, 63 (2021).

[30] Kovacs, J. A., Chacón, P. & Abagyan, R. Predictions of protein flexibility: First-order measures. Proteins: Structure, Function, and Bioinformatics 56, 661–668 (2004).

[31] Bahar, I., Lezon, T. R., Bakan, A. & Shrivastava, I. H. Normal Mode Analysis of Biomolecular Structures: Functional Mechanisms of Membrane Proteins. Chemical Reviews 110, 1463–1497 (2010).

[32] Dobbins, S. E., Lesk, V. I. & Sternberg, M. J. E. Insights into protein flexibility: The relationship between normal modes and conformational change upon protein–protein docking. Proceedings of the National Academy of Sciences 105, 10390–10395 (2008).

[33] Fuglebakk, E., Tiwari, S. P. & Reuter, N. Comparing the intrinsic dynamics of multiple protein structures using elastic network models. Biochimica et Biophysica Acta (BBA) - General Subjects 1850, 911–922 (2015).

[34] Balog, E. et al. Direct Determination of Vibrational Density of States Change on Ligand Binding to a Protein. PHYSICAL REVIEW LETTERS 93, 4 (2004).

[35] Echave, J. Why are the low-energy protein normal modes evolutionarily conserved? Pure and Applied Chemistry 84, 1931–1937 (2012).

[36] Micheletti, C. Comparing proteins by their internal dynamics: exploring structure-function relationships beyond static structural alignments. Physics of Life Reviews 10, 1–26 (2013).

[37] Zhang, S., Li, H., Krieger, J. M. & Bahar, I. Shared Signature Dynamics Tempered by Local Fluctuations Enables Fold Adaptability and Specificity. Molecular Biology and Evolution 36, 2053–2068 (2019).

[38] De Geyter, J. et al. Trigger factor is a bona fide secretory pathway chaperone that interacts with SecB and the translocase. EMBO reports 21 (2020).

[39] Tsirigotaki, A., Elzen, R. V., Veken, P. V. D., Lambeir, A.-M. & Economou, A. Dynamics and ligand-induced conformational changes in human prolyl oligopeptidase analyzed by hydrogen/deuterium exchange mass spectrometry. Scientific Reports 7, 2456 (2017).

[40] Bai, Y., Milne, J. S., Mayne, L. & Englander, S. W. Primary structure effects on peptide group hydrogen exchange. Proteins: Structure, Function, and Bioinformatics 17, 75–86 (1993).

[41] Connelly, G. P., Bai, Y., Jeng, M.-F. & Englander, S. W. Isotope effects in peptide group hydrogen exchange. Proteins: Structure, Function, and Genetics 17, 87–92 (1993).

[42] Mori, S., Zijl, P. C. M. v. & Shortle, D. Measurement of water–amide proton exchange rates in the denatured state of staphylococcal nuclease by a magnetization transfer technique. Proteins: Structure, Function, and Bioinformatics 28, 325–332 (1997).

[43] Nguyen, D., Mayne, L., Phillips, M. C. & Walter Englander, S. Reference Parameters for Protein Hydrogen Exchange Rates. Journal of the American Society for Mass Spectrometry 29, 1936–1939 (2018).

[44] Zhang, Z. & Smith, D. L. Determination of amide hydrogen exchange by mass spectrometry: a new tool for protein structure elucidation. Protein Science: A Publication of the Protein Society 2, 522–531 (1993).

[45] Hvidt, A. & Nielsen, S. O. Hydrogen Exchange in Proteins. In Anfinsen, C. B., Anson, M. L., Edsall, J. T. & Richards, F. M. (eds.) Advances in Protein Chemistry, vol. 21, 287–386 (Academic Press, 1966).

[46] Harris, C. R. et al. Array programming with NumPy. Nature 585, 357–362 (2020).

[47] team, T. p. d. pandas-dev/pandas: Pandas (2020). URL https://doi.org/10.5281/zenodo.3509134.

[48] Virtanen, P. et al. SciPy 1.0: Fundamental Algorithms for Scientific Computing in Python. Nature Methods (2020).

[49] Walt, S. v. d. et al. scikit-image: image processing in Python. PeerJ 2, e453 (2014).

[50] Roelfs, M. & Kroon, P. C. tBuLi/symfit: symfit 0.5.3 (2020). URL https://zenodo.org/record/4020630.

[51] Paszke, A. et al. PyTorch: An Imperative Style, High-Performance Deep Learning Library. In Wallach, H. et al. (eds.) Advances in Neural Information Processing Systems 32, 8024–8035 (Curran Associates, Inc., 2019).

[52] Dask Development Team. Dask: Library for dynamic task scheduling (2016).

[53] Smit, J. Jhsmit/HDXrate: Zenodo release (2021). URL https://zenodo.org/record/4630881.

[54] Hunter, J. D. Matplotlib: A 2D graphics environment. Computing in science & engineering 9, 90–95 (2007).

[55] Bokeh Development Team. Bokeh: Python library for interactive visualization (2020).

[56] Kluyver, T. et al. Jupyter Notebooks – a publishing format for reproducible computational workflows. In Loizides, F. & Scmidt, B. (eds.) Positioning and Power in Academic Publishing: Players, Agents and Agendas, 87–90 (IOS Press, 2016). URL https://eprints.soton.ac.uk/403913/.

[57] Rudiger, P. et al. holoviz/panel: Version 0.11.3 (2021). URL https://zenodo.org/record/4692827.

[58] Rose, A. S. & Hildebrand, P. W. NGL Viewer: a web application for molecular visualization. Nucleic Acids Research 43, W576–W579 (2015).

[59] Rose, A. S. et al. NGL viewer: web-based molecular graphics for large complexes. Bioinformatics 34, 3755–3758 (2018).

[60] Tiwari, S. P. et al. WEBnm@ v2.0: Web server and services for comparing protein flexibility. BMC Bioinformatics 15, 427 (2014).

## References

[1] Guillet, V. et al. Structural insights into chaperone addiction of toxin-antitoxin systems. Nature Communications 10, 782 (2019).

[2] F, M. et al. The EMBL-EBI search and sequence analysis tools APIs in 2019. Nucleic Acids Research 47, W636–W641 (2019).

[3] Tiwari, S. P. et al. WEBnm@ v2.0: Web server and services for comparing protein flexibility. BMC Bioinformatics 15, 427 (2014).

[4] Dobbins, S. E., Lesk, V. I. & Sternberg, M. J. E. Insights into protein flexibility: The relationship between normal modes and conformational change upon protein-protein docking. Proceedings of the National Academy of Sciences 105, 10390–10395 (2008).

[5] Ferbitz, L. et al. Trigger factor in complex with the ribosome forms a molecular cradle for nascent proteins. Nature 431, 590–596 (2004).

[6] Morgado, L., Burmann, B. M., Sharpe, T., Mazur, A. & Hiller, S. The dynamic dimer structure of the chaperone Trigger Factor. Nature Communications 8, 1992 (2017).

[7] Ku, B., Liang, C., Jung, J. U. & Oh, B.-H. Evidence that inhibition of BAX activation by BCL-2 involves its tight and preferential interaction with the BH3 domain of BAX. Cell Research 21, 627–641 (2011).

[8] Petros, A. M. et al. Solution structure of the antiapoptotic protein bcl-2. Proceedings of the National Academy of Sciences 98, 3012–3017 (2001).

[9] Szeltner, Z., Rea, D., Renner, V., Fülöp, V. & Polgár, L. Electrostatic Effects and Binding Determinants in the Catalysis of Prolyl Oligopeptidase: SITE-SPECIFIC MUTAGENESIS AT THE OXYANION BINDING SITE *. Journal of Biological Chemistry 277, 42613–42622 (2002).

[10] Yuan, B. et al. Functional self-nonamerization of the Type III translocase chaperone/exported protein receptor. bioRxiv 2020.11.20.391094 (2020).

[11] Dekker, C., Kruijff, B. d. & Gros, P. Crystal structure of SecB from Escherichia coli. Journal of Structural Biology 144, 313–319 (2003).

[12] Gelis, I. et al. Structural Basis for Signal-Sequence Recognition by the Translocase Motor SecA as Determined by NMR. Cell 131, 756–769 (2007).

[13] Sharma, V. et al. Crystal structure of Mycobacterium tuberculosis SecA, a preprotein translocating ATPase. Proceedings of the National Academy of Sciences 100, 2243–2248 (2003).

[14] Konno, M., Ito, M., Hayano, T. & Nobuhiro. The Substrate-binding Site inEscherichia coliCyclophilin A Preferably Recognizes acis-proline Isomer or a Highly Distorted Form of thetransIsomer. Journal of Molecular Biology 256, 897–908 (1996).

[15] Konno, M. et al. Escherichia coli cyclophilin B binds a highly distorted form of trans-prolyl peptide isomer. European Journal of Biochemistry 271, 3794–3803 (2004).

[16] Quiocho, F. A., Spurlino, J. C. & Rodseth, L. E. Extensive features of tight oligosaccharide binding revealed in high-resolution structures of the maltodextrin transport/chemosensory receptor. Structure 5, 997–1015 (1997).

